# Genomic characterization of *Escherichia coli* harbor a polyketide synthase (*pks*) island associated with colorectal cancer (CRC) development

**DOI:** 10.1101/2024.06.16.599199

**Authors:** Chao Lv, Mohd Abdullah, Weiye Chen, Nan Zhou, Zile Cheng, Yiwen Chen, Min Li, Kenneth W. Simpson, Ahmed Elsaadi, Yongzhang Zhu, Steven M. Lipkin, Yung-Fu Chang

## Abstract

The *E. coli* strain harboring the polyketide synthase (*Pks)* island encodes the genotoxin colibactin, a secondary metabolite reported to have severe implications for human health and for the progression of colorectal cancer. The present study involved whole-genome-wide comparison and phylogenetic analysis of *pks* harboring *E. coli* isolates to gain insight into the distribution and evolution of these organism. Fifteen *E. coli* strains isolated from patients with ulcerative colitis were sequenced, 13 of which harbored pks islands. In addition, 2,654 genomes from the public database were also screened for *pks* harboring *E. coli* genomes, 158 of which were *pks*-positive isolates. Whole-genome-wide comparison and phylogenetic analysis revealed that 171 (158+13) *pks*-positive isolates belonged to phylogroup B2, and most of the isolates associated to sequence types ST73 and ST95. One isolate from an ulcerative colitis (UC) patient was of the sequence type ST8303. The maximum likelihood tree based on the core genome of *pks*-positive isolates revealed horizontal gene transfer across sequence types and serotypes. Virulome and resistome analyses revealed the preponderance of virulence genes and a reduced number of antimicrobial genes in *Pks*-positive isolates. This study strongly contributes to understanding the evolution of *pks* islands in *E. coli*.

## Introduction

*Escherichia coli* (*E. coli*) is a pathogenic microbe that inhabits the gut of animals and humans and accounts for intestinal and extraintestinal infections[1, 2]. The diverse population of *E. coli* is widely distributed in eight major phylogenetic groups (A, B1, B2, C, D, E, F, and G) [3]. A major population of *E. coli* belongs to phylogroup B2, which causes severe infections such as urinary tract infection (UTI), sepsis, pneumonia, and neonatal meningitis [4]. There are multiple reasons for the evolution of virulence in *E. coli*, but a major role is played by horizontal gene transfer, point mutation, and inactivation of antivirulence genes [5, 6]. Virulence factors such as toxins, adhesins, capsules, and iron acquisition systems are often encoded by genes that can be mobilized through various methods, including mobile genetic elements, genomic islands, phages, and plasmids. Horizontal gene transfer allows the widespread distribution of these genes in extraintestinal pathogenic *E. coli* (ExPEC) strains [7, 8]. Genomic islands, large regions of more than 10 kb that are often bounded by repetitive structures and carry mobility factors such as integrase and transposes, exhibit associations with tRNA genes and have diverse G+C contents [9]. Pathogenicity islands (PAIs), a small subgroup of genomic islands, play pivotal roles in the evolution of bacterial virulence by incorporating virulence-associated factors and adaptive horizontal gene transfer [10, 11]. PAI encodes a toxin known as colibactin, a nonribosomal peptide-polyketide secondary metabolite and is observed in strains of *E. coli* associated with uropathogenic, commensal, and neonatal meningitis [12]. Colibactin can induce double-stranded DNA breaks in eukaryotic cells, leading to cell cycle arrest at the G_2_-M phase and chromosomal aberrations [13, 14]. It significantly contributes to severe clinical manifestations such as meningitis [15] and sepsis [12].

Colibactin, known for its virulence against the extraintestinal pathogen *E. coli* (ExPEC), is also suspected of its role as a procarcinogen factor [14, 16, 17]. Elevated levels of harmful *E. coli* strains are frequently detected in CRC patients compared to healthy individuals [18]. Several recent studies suggest that certain strains of *E. coli* that possess the *pks* Island may play a causal role in the development of human CRC [17, 19, 20]. The colibactin pks+ gene is significantly enriched in CRC patients, in familial adenomatous polyposis (FAP), and in DNA mismatch repair-deficient CRC patients [21, 22]. According to preclinical studies, pks+ *E. coli* drives tumorigenesis and increases tumor burden in several CRC and FAP mouse models [21, 23, 24]. Notably, colibactin-induced DNA damage creates a specific mutational signature in CRC tumors that can be computationally monitored and used to measure the contribution of pks+ *E. coli* to tumor burden[25, 26].

The biosynthesis machinery of colibactin is located on the *PKS* island, spanning a region of 54 kb and housing 19 genes. These genes included nonribosomal peptide mega synthases (NRPSs; clbH, clbJ, and clbN), polyketide mega synthases (*PKSs*; clbC, clbI, and clbO), two hybrid NRPS-PKSs (clbB and clbK), and nine accessory and tailoring enzymes [13]. The presence of the *pks* island is not confined to pathogenic organisms; it has also been observed in commensal and probiotic bacterial strains [27]. Its presence extends beyond *E. coli*, encompassing members of the Enterobacteriaceae family, such as *Citrobacter koseri*, *Klebsiella pneumoniae*, and *K. aerogenes* [28]. The association between *pks-*positive (*pks^+^*) *E. coli* and colorectal cancer is evident in biopsy samples, revealing an elevated prevalence of *pks+* islands harboring *E. coli* [17, 29]. Notably, these isolates are found in more than half of patients with familial adenomatous polyps and contribute to carcinogenesis through mucus degradation, adherence, and enhanced colonization within colonic biofilms[30]. In addition to their speculated role in CRC progression, *pks* islands serve as virulence factors with clinical implications, contributing to systemic infection, neonatal meningitis, and lymphopenia, according to various studies[31–33].

In this study, we performed whole-genome sequencing (WGS) of 15 *E. coli* isolates from patients with ulcerative colitis (UC) and performed genome-wide comparisons and phylogenetic analysis of *pks* islands harboring *E. coli* isolates from 15 UC strains and 2654 datasets from the NCBI database. The study describes the distribution of *pks^+^ E. coli* among phylogroups, STs, and serogroups, followed by core and pangenome analysis. A phylogenomic study is also being performed on the core genome to understand island acquisition and evolution. The antibiotic resistance genes and virulence genes were mined to understand the drug resistance and virulence characteristics of *pks* harboring *E. coli* isolates.

## Materials and methods

### Whole-genome sequencing of *E. coli*

A total of 15 *E. coli* strains from separate patients with ulcerative colitis (UC) were used in this study, and the strains were previously reported in 2004 [34]. The genomic DNA of the 15 strains was extracted using the TIANamp Bacteria D.N.A. Kit (TIANGEN, Beijing, China). Subsequently, the DNA libraries were prepared using the KAPA HyperPrep Kit (Roche, Basel, Switzerland) following the manufacturer’s instructions and sequenced on the Illumina NovaSeq platform with a 150 bp paired-end strategy. Furthermore, the strain HM229 was subjected to long-read sequencing using an Oxford Nanopore Technology (ONT.) MinION device.

The draft genomes were assembled using the PGCGAP pipeline with the SPAdes v3.13.1 algorithm, and the long-read genomes were assembled using the Unicycler v0.5.0 algorithm (https://gitee.com/liaochenlanruo/pgcgap) [35]. To explore the genomic characteristics of *E. coli* isolates harboring polyketide synthase (*pks*) islands from patients with UC, an additional dataset of 2654 complete genomes of *E. coli* was downloaded from the NCBI (deadline 2022.12.31, **Table S1)**. Moreover, the strains in the downloaded dataset included information on the host, host disease, geographic location, and collection date.

### Identification of *pks^+^* genes in *E. coli*

The reference sequence of the *pks+* island (GenBank accession number: AM229678.1) was downloaded from NCBI and used as a query file to perform BLASTn against the genomes of *E. coli* strains from UC patients and all downloaded genomes. The identity and query coverage thresholds of BLASTn were 85%.

### Phylogenetic analysis of *E. coli* strains

The phylogroups of the *E. coli* genomes (2654 downloaded strains and 15 strains from UC patients) were detected using ClermonTyping (https://github.com/A-BN/ClermonTyping) [36]. Sequence typing (ST) was performed using MLST (https://github.com/tseemann/mlst) [37]. ECTyper (https://github.com/phac-nml/ecoli_serotyping) was used to perform serotyping of the genomes [38]. Subsequently, the *pks*^+^ strains were filtered, and the minimum spanning tree was generated based on the STs in PHYLOViZ 2.0 (https://www.phyloviz.net/) using the goeBURST algorithm [39]. After annotation by Prokka (http://vicbioinformatics.com/)[40], the software Roary 3.11.2 (http://sanger-pathogens.github.io/Roary/) [41] was used to determine the core genes of the *pks*^+^ strains. The genomic core gene-based phylogenetic tree was subsequently constructed using FastTree 2.1 (http://meta.microbesonline.org/fasttree/) [42], which infers an approximately maximum likelihood algorithm with generalized time-reversible (GTR) models. The core-gene-based phylogenetic tree was visualized using Interactive Tree Of Life (iTOL, https://itol.embl.de/).

To explore the divergence time of the strains harboring *pks* islands from UC patients, we tracked the collection time of each *pks*^+^ isolate in the NCBI database. The collection time of strains from UC patients was defined in 2004. The core gene-based phylogenetic tree was subsequently constructed as described above, and Bayesian inference of divergence dates on the bacterial phylogenetic trees was implemented via an R package called BactDating, which is freely available for download at https://github.com/xavierdidelot/BactDating [43]. The results were displayed in Figtree v1.4.4 (http://tree.bio.ed.ac.uk/software/figtree/).

### Virulome and resistome profiling of *pks*^+^ strains from UC patients

To explore the characteristics of the virulome and resistome of the *pks*^+^ strains from UC patients, a total of 102 isolates (including *pks^+^* and *pks*^-^; **Table S2**) were selected from among the 2669 strains following the established standards: (1) Based on the STs of UC patient strains, 10 *pks*^-^ and *pks*^+^ strains were selected randomly if the STs had more than 10 *pks*^-^ and *pks*^+^ strains, similar to ST95; (2) Ten *pks*^-^ and *pks*^+^ strains were selected if the STs only had more than 10 *pks*positive or negative strains, such as ST127, ST73, ST453, and ST131; (3) The strains were all selected if the STs had fewer than 10 strains, similar to ST141.

Then, 102 genomic core gene-based phylogenetic trees were constructed as above to display the phylogeny of the *pks*^-^ and *pks*^+^ strains. All the assemblies were screened for antimicrobial resistance genes (ARGs) and virulence genes (VGs) against the comprehensive Antibiotic Resistance Database (CARD) [44] and Virulence Factor Database (VFDB) [45] using Abricate (https://github.com/tseemann/abricate). The numbers of ARGs and VGs in various comparison groups (*pks*^+^ strains from UC patients versus (*VS*) *pks*^+^ strains from others and *pks*^-^ strains; and *pks*^+^ strains from UC patients *vs. pks*^+^ strains from urinary tract infection (UTI) patients (10) and healthy people (11); **Table S1**) were visualized using boxplots generated with ggplot2 v3.3.2 in R. Meanwhile, the percentages of ARGs and VGs in various comparison groups were visualized using a dot plot in ggplot2 v3.3.2 in R.

### Chromosomal and plasmid maps of the UC isolate HM229

A chromosomal map of strain HM229 (phylogroup B, ST12) was created via long-read sequencing and compared to those of 7 other *pks*^+^ strains randomly selected from each phylogroup, including the reference strain IHE3034 (**Table S3**), utilizing the BLAST Ring Image Generator (BRIG, https://sourceforge.net/p/brig). The annotated chromosome of HM229 was used as a reference for generating whole chromosomal sequence comparisons. The plasmid sequences of the 8 strains were filtered out, and comparative plasmid maps were generated using BRIG. Moreover, the genes on the plasmid of HM229 were retrieved from the UniProtKB database (https://doi.org/10.1093/nar/gkac1052). Additionally, the location of putative phage or phage-like regions of the 8 strains was determined using PHASTER (https://phaster.ca/) [46, 47]. Genomic islands (GIs) in the 8 strains were predicted using the online tool IslandViewer (https://www.pathogenomics.sfu.ca/islandviewer) [48]. Since multiple methods are integrated by this online prediction when determining the start and end positions of GI, the overlapping regions were combined. The predicted phage or phage-like regions and GIs were labeled on the chromosomal maps.

### Whole-genome alignment of eight *pks^+^* strains

A multiple genome alignment tool called Mauve (https://darlinglab.org/mauve/user-guide/screenshots.html) was used to construct and visualize the whole chromosomal alignment of the selected 8 strains. Mauve compares multiple genome sequences and finds regions of homology called locally collinear blocks (LCBs). The progressive Mauve algorithm was used with the default parameters.

### Analysis *of the pks* island structure

The HM229 strain was compared with the other 7 selected strains (**Table S3**), which were attributed to different phylogroups according to *pks* island structure analysis, and the IHE3034 strain was used as a reference. First, the *pks* island sequence was extracted from the whole genome and annotated by Prokka (http://vicbioinformatics.com/) to accrue the GBK format of the *pks* island sequence. The GBK files of the 8 strains were subsequently submitted to the software Easyfig2.2.5 (https://mjsull.github.io/Easyfig/) to create linear comparison figures of multiple genomic loci with an easy-to-use graphical user interface (GUI) [49].

### SNP analysis of whole genomes and *pks* genes

For SNP analysis, the 7 selected strains were mapped to the genome of HM229 by the snippy program (https://github.com/tseemann/snippy). The recombinant region was removed from the resulting alignment by the Gubbins program, and then core SNPs were extracted by the SNP-sites program. In addition to the SNP analysis of the whole genome of the 8 strains, the sequence of the *pks* island was also extracted after BLASTn to conduct SNP analysis.

### Statistical analysis

Categorical data were analyzed via the chi-square test. Continuous data with normal or nonnormal distributions were analyzed with a *t* test or Mann □ Whitney U test. For comparisons of multiple groups, analysis of variance (ANOVA) or Kruskal□ Wallis H test was used. All the statistical analyses were performed in IBM SPSS Statistics 25 (IBM, Armonk, USA).

## Results

### Genomic analysis revealed *pks* island in *E. coli* strains

Blastn revealed that 13 of the 15 (86.67%) *E. coli* strains from UC patients harbored *pks* island, and the number of *pks^+^* strains was 158 (158/2654, 5.95%) according to the downloaded genomic dataset from the NCBI (**Table S1**); these strains included strains collected from humans (8.98%, 103/1147), food animals (2.07%, 9/435), laboratories (8.99%, 8/89), wildlife (3.85%, 7/182), environmental samples (2.37%, 5/211), companion animals (6.67%, 4/60), food samples (0.82%, 1/122), marine organisms (14.29%, 1/7) and undefined sources (4.99%, 20/401). Moreover, the *pks^+^* percentage of strains from patients with urinary tract infection (10/69, 14.49%), bacteremia (5/68, 7.35%), diarrhea (1/63, 1.59%), sepsis (2/26, 7.69%), cystitis (2/14, 14.29%), gastroenteritis (0/14, 0.00%), hemolytic uremic syndrome (0/10, 0.00%), and healthy status (11/69, 15.94%) were significantly lower than those from UC patients **(Table 1).**

**Table 1:**
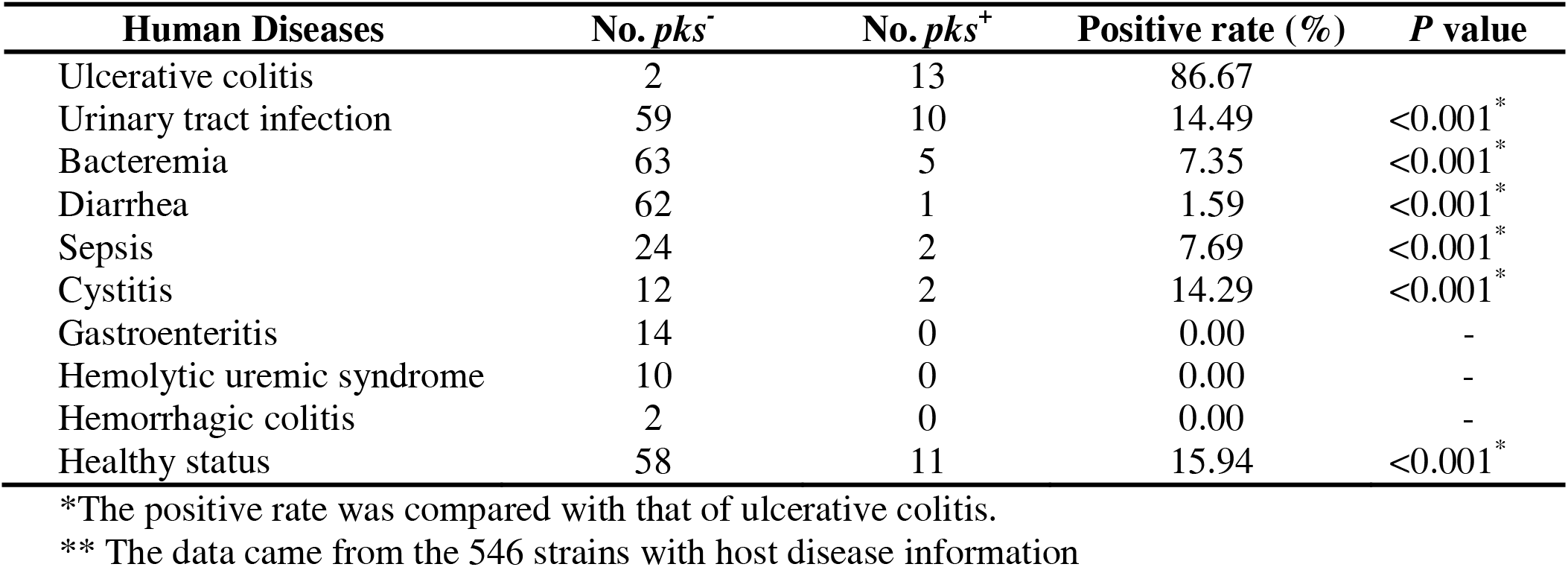
The occurrence of *pks* island in *E. coli* strains from humans with different diseases**.

### Distribution of *pks^+^ E. coli*

The phylogroup analysis showed that the 13 *pks^+^* strains from UC patients belonged to phylogroup B2 (100%), and the absolute dominant occurrence of *pks* island in phylogroup B2 also appeared in 158 downloaded genomes (95.57%, 151/158; **Table 2**). The 13 *pks^+^* strains from U.C. patients belonged to 6 different STs, namely, ST12, ST73, ST95, ST127, ST141, and ST8303 **(Table 2),** and the strains belonging to ST12 (30.77%, 4/13) had the highest percentage of occurrence. Among the 158 downloaded *pks*^+^ strains, ST73 (29.75%, 47/158), ST95 (22.78%, 36/158) and ST127 (15.19%, 24/158) were the dominant STs. Furthermore, compared with the STs of the downloaded *pks*^+^ genomes, the ST8303 sequence was distinct. The minimum spanning tree showed that the 13 *pks^+^* isolates of UC patients were mainly assigned to three clonal complexes (CC) of CC12, CC95, and CC73, similar to the 158 downloaded *pks*^+^ genomes **(Fig. 1).**

**Fig. 1.**
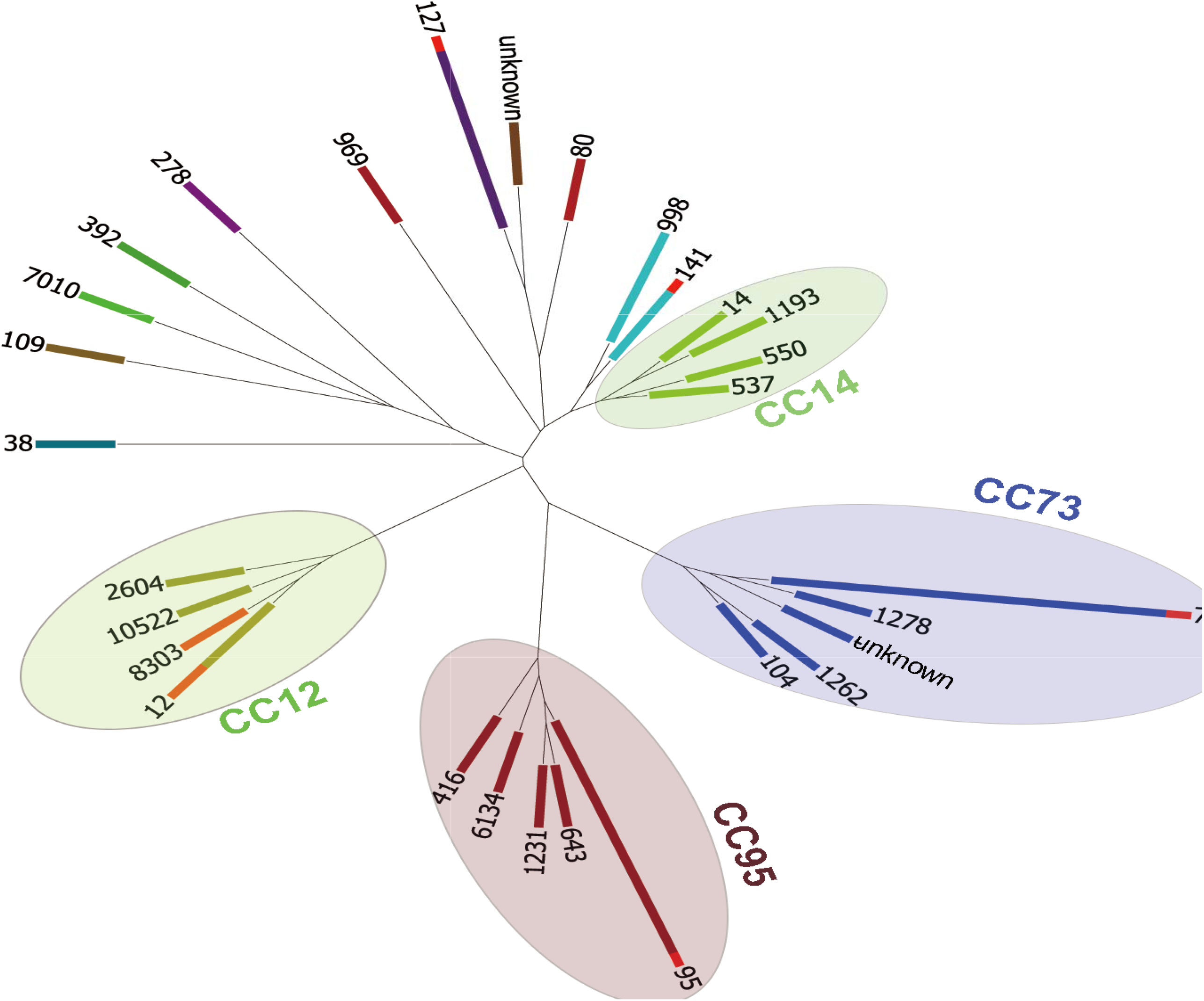
The minimum spanning tree based on the STs of all *pks*^+^ strains (n=171). The four colored shadow circles represent different CCs, the strip bar represents the number of strains, and the difference between some strip bars represents the percentage of strains from UC patients.

**Table 2:**
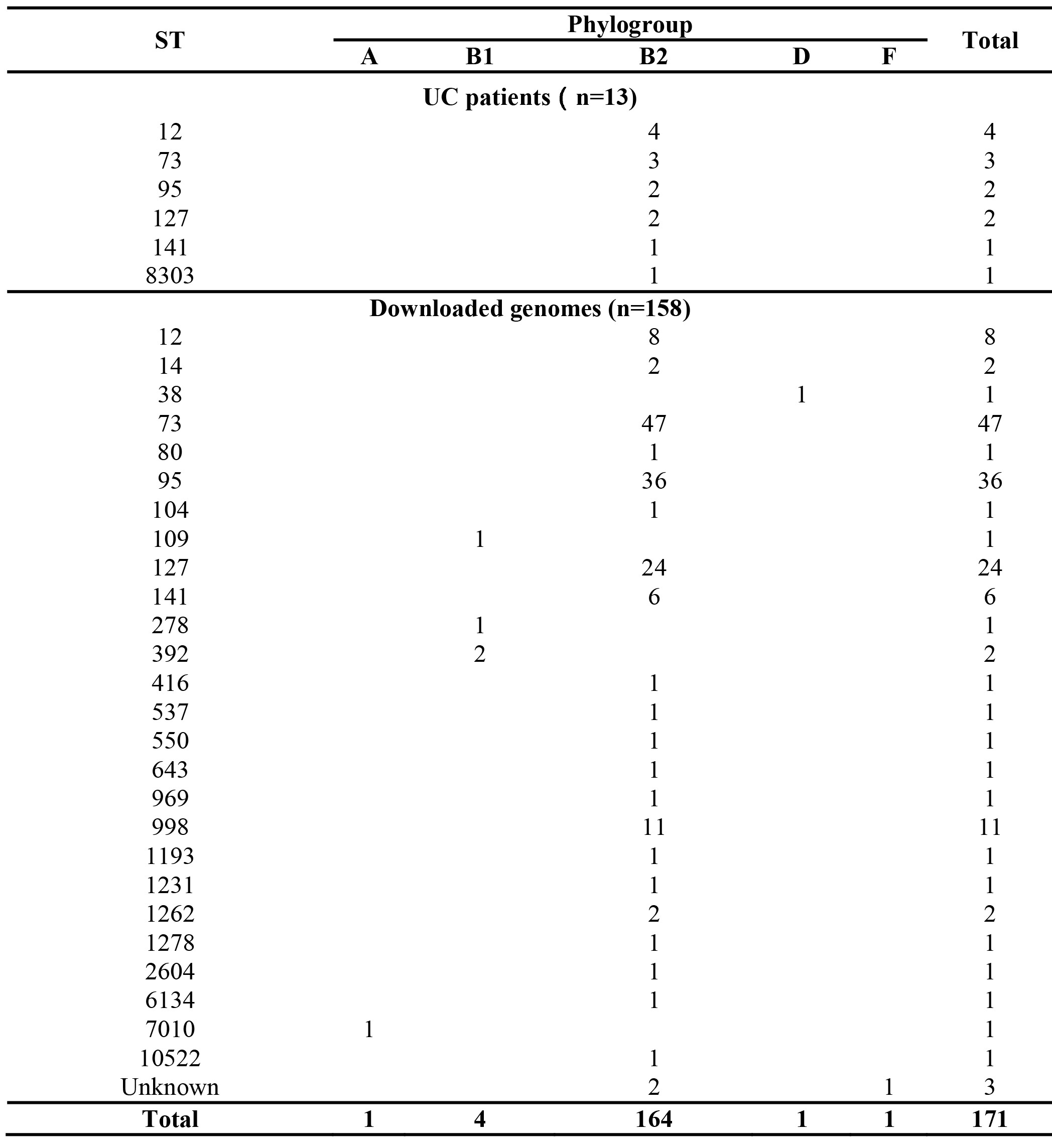
Distribution of the *pks^+^* strains according to sequence type (ST.)

There are 10 serotypes in the strains of UC patients, and there are no dominant serotypes. Among the 158 downloaded strains, O6:H1 (25.95%, 41/158), O6:H31 (13.92%, 22/158), and O18:H7 (13.92%, 22/158) were the three main dominant serotypes **(Table S2, Fig. 2).** Additionally, from the phylogenetic tree, the predominant serotype in CC73 was O6:H1 (74.55%, 41/55), the dominant serotype in CC95 was O18:H7 (51.22%, 21/41), and that in CC12 was O4:H5 (76.92%, 10/13).

**Fig. 2.**
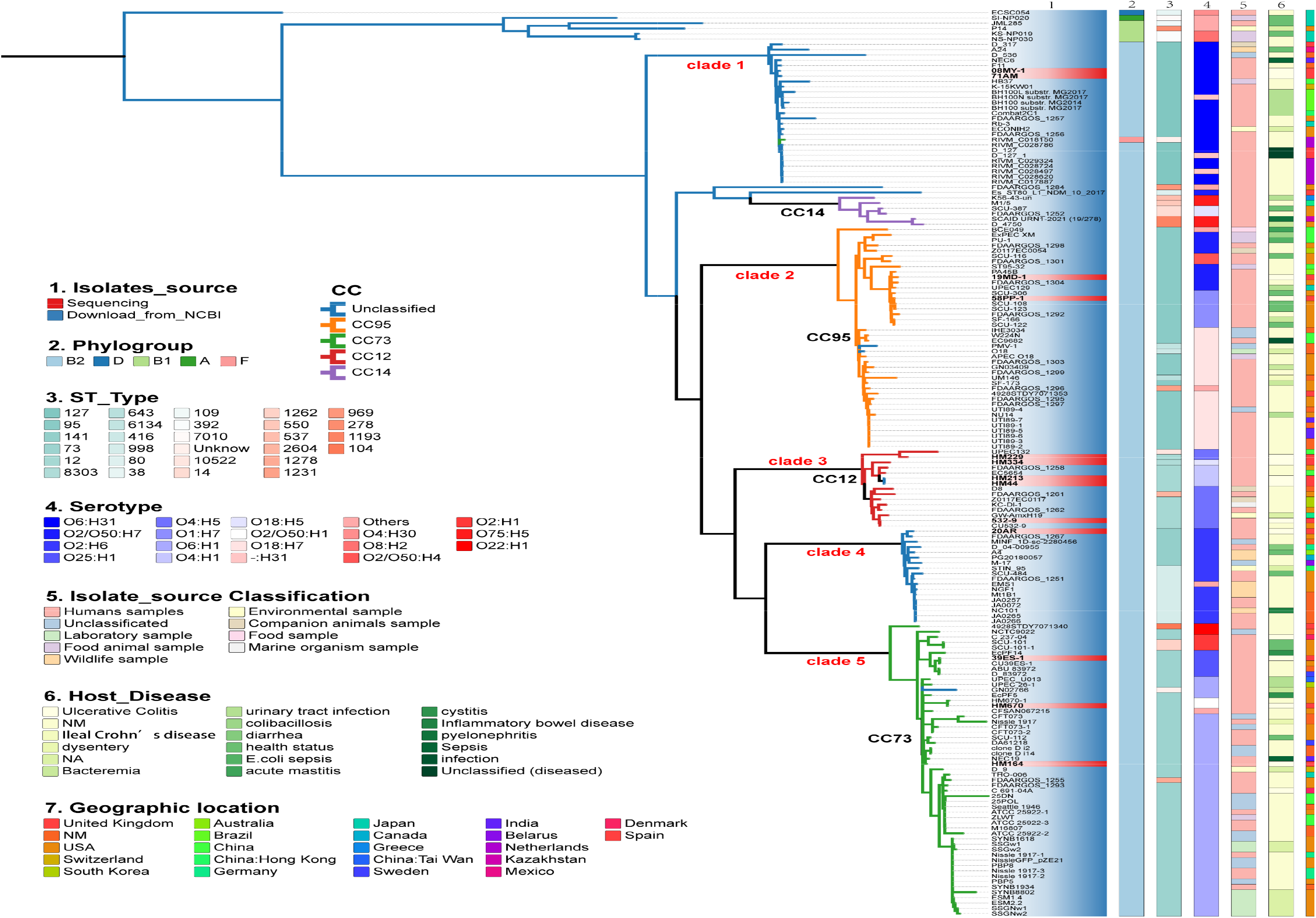
Phylogenetic inference based on the analysis of core genes of 171 *pks^+^* strains. The core gene-based ML tree was constructed based on 3482 core genes. NM (column 6): not mentioned; NA: not applicable.

### Phylogenetic analysis of *pks^+^ E. coli* and their divergence time

The core genome maximum-likelihood phylogeny was obtained from Fast-Tree.C. The 13 pks+ strains from UC patients were distributed into 5 different clades (we named clades 1-5). Five strains (5/13) were assigned to clade 3, which was the dominant branch of CC12 **(Fig. 2).** Interestingly, the clades of the 13 isolates identified by core-gene-based phylogeny were strict, with the clusters shown in the minimum spanning tree based on the STs. Among the five clades, the strains from humans made up a dominant proportion of the strains, and the strains from the laboratory were mainly attributed to clade 5 (CC73), the strains from wildlife were mostly attributed to clade 4, and the strains from companion animals were primarily attributed to clade 3 (CC12). For the serotype, O6:H31, O18:H17, O4:H5, O2:H6, and O6:H1 were the dominant serotype from clade 1 to clade 5, respectively (**Fig. 2**). However, 2 of the 3 *pks^+^* isolates from UC patients belonged to clade 5 (CC73), which was another two serotypes (O25:H1, O2/O50:H1); moreover, the UC patient-derived *pks^+^* strains were not the dominant serotype in clade 2 (CC12) or 3 (CC95) **(Fig. 2).**

The core gene-based phylogenetic tree with divergence time showed that the isolate coded HM670, which belongs to ST73, had the earliest divergence time among the 13 strains from UC patients, indicating that the acquisition of the *pks* island first occurred in UC *E. coli* strains from ST73. Moreover, the strains coded HM213 and HM44, which are ST12 strains, had a divergence time of 26.23 (**Fig. 3**). For the three main CCs of the UC *E. coli* strains, CC73 had the earliest average divergence time of 91.97, and CC95 had the latest average divergence time of 34.48; these findings are the same as the comparison results for all the *pks^+^* strains, which were displayed by the configuration of the phylogenetic tree. Moreover, the divergence time of phylogroup B2 occurred earlier than that of the other phylogroups A, B1, D, and F, supporting the hypothesis that the *pks* island first occurred in the *E. coli* belonging to phylogroup B2. Interestingly, the *pks^+^ E. coli* strains pertained to phylogroups A, B1, and D formed a divergent cluster (**Fig. 3**). We also compared the divergence times of *pks^+^ E. coli* strains from urinary tract infection (UTI), bacteraemia, and health status. In ST73, the earliest divergence time was from healthy people, followed by that of UC patients, and the shortest was that of UTI. In ST127, the order of divergence time was U.T.I. (74.67)> healthy people (67.74) > U.C. (43.46). In contrast, in ST95, strains from bacteremia had the earliest divergence time (67.89), followed by strains from UTI (49.49) and UC (39.23), and the latest diverged from healthy people (29.63, **Fig. 3**).

**Fig. 3.**
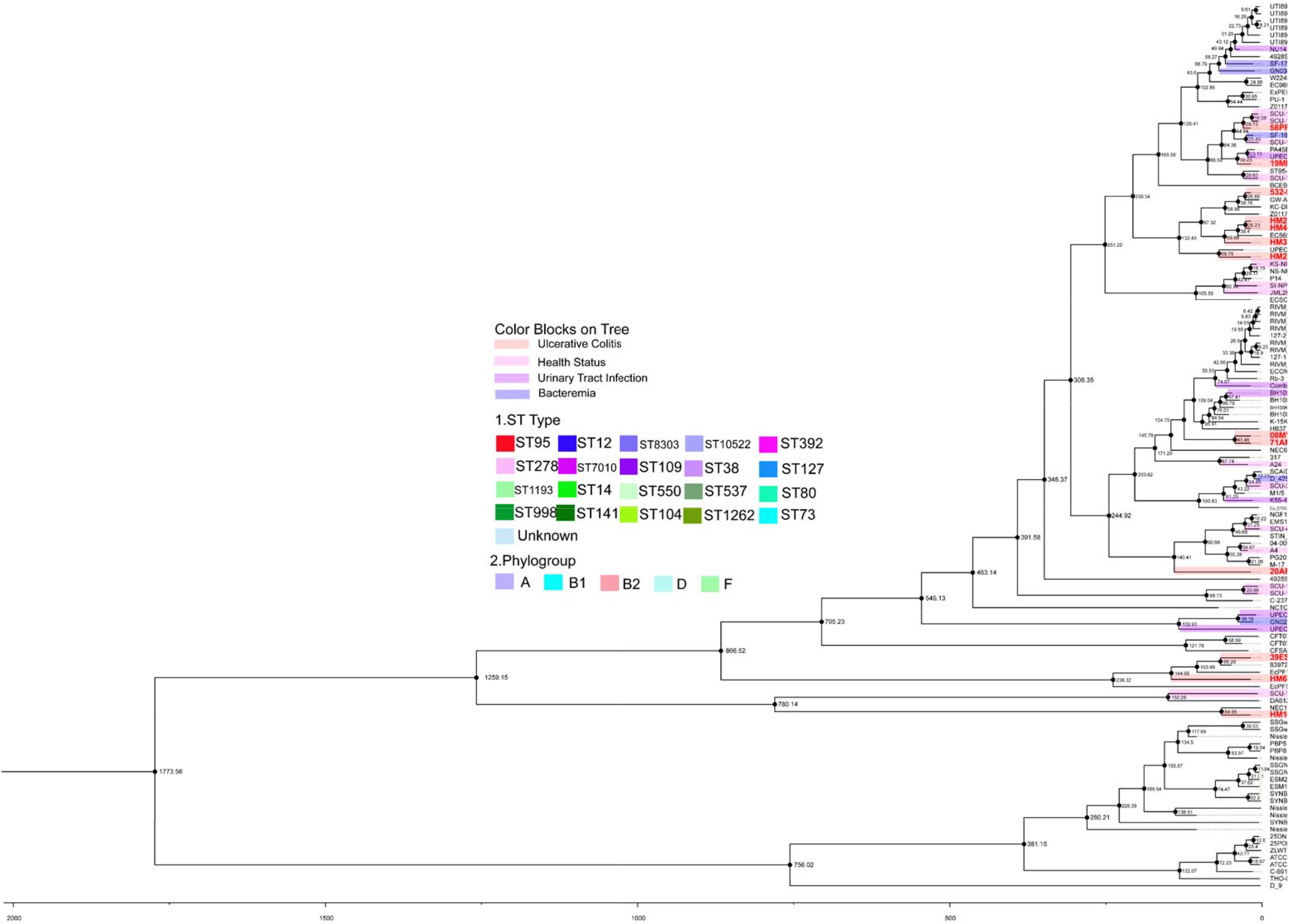
The core gene phylogenetic tree with a divergence time of 124 *pks*^+^ strains.

### Pangenome analysis of *pks^+^ E. coli* strains

The median number of core genes associated with the *pks^+^* strains from UC patients was 3470.00 (IQR: 3467.00-3473.00), which was slightly lower than that associated with the *pks^+^* strains downloaded from NCBI (3475.00, IQR: 3466.00-3479.00, *P*=0.088; Mann–Whitney U test; **Fig. S1A**). However, the median number of accessory genes associated with the *pks^+^* strains from UC patients (1333.00, IQR:1185.00-1395.00) was greater than that associated with the *pks^+^* strains downloaded from NCBI (1316.00, IQR: 1191.00-1453.00), and the distribution was also not different (*P*=0.877, Mann–Whitney U test; **Fig. S1B**).

The 54 kb *pks* island contains 19 genes (*clb*A to-S) encoding biosynthetic machinery. A heatmap of the pangenome analysis revealed four gene deletions, namely, *clb*J, *clb*H, *clb*M and *clb*I, in the *pks* island region of strains from UC patients (except for the HM229 strain) compared with most *pks^+^* isolates and the reference isolate IHE3034 (**Fig. 4**). Additionally, the UC patient-derived strain 532-9 had two additional deletions of *clb*B and the putative transposase gene. According to previous research, the *pks* island genes are involved in enzymatic interactions, and *clb*J, *clb*H, *clb*I, and *clb*B are involved in the assembly of mega synthase nonribosomal peptide synthetase (NRPS) and polyketide synthase (PKS) genes, while the *clb*M gene effluxes precolibactin, which is unloaded from the aforementioned assembly line through the periplasm.

**Fig. 4.**
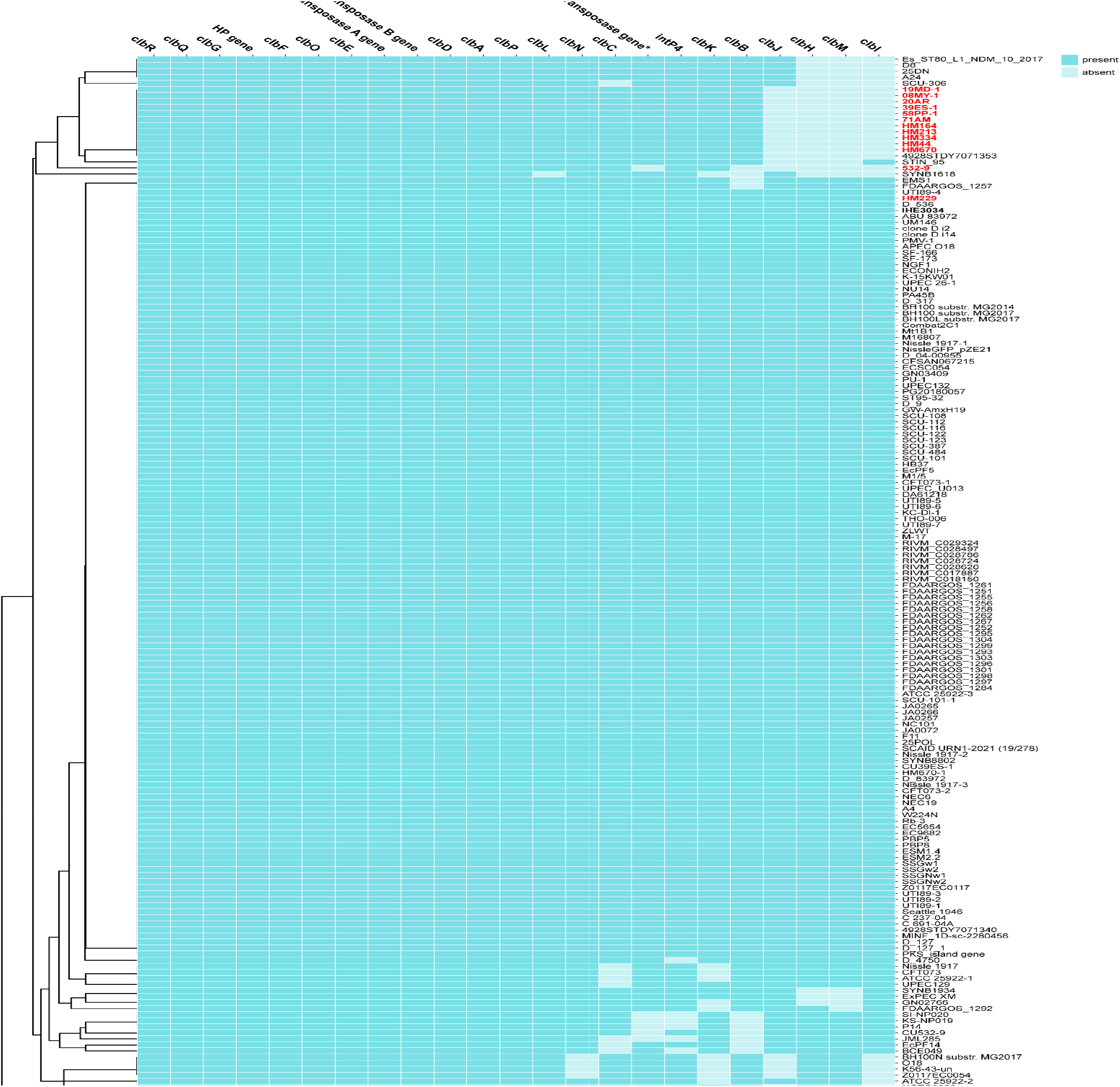
The heatmap of the presence or absence of the genes of the *pks* island filter from the 171 *pks*^+^ *E. coli* isolates. The red color of the isolates name indicates the isolates from UC patients. The method of heatmap clustering is based on the complete Euclidean distance.

### Resistome and Virulome analysis of *pks^+^ E. coli* strains

From the phylogenetic tree of the 102 selected *E. coli* strains, the *pks^+^* or *pks^-^* strains exhibited an obvious clustering pattern (**Fig. 5A, D**), despite the strains belonging to ST95, ST12, and ST141 containing both *pks^+^* or *pks^-^* strains. Interestingly, of the 2669 strains, the strains belonging to ST73 (n=50), ST127 (n=26), and ST998 (n=11) all harbored *pks* islands **(Table S3)**. The resistome analysis revealed that the maximum quantity of ARGs among the 102 selected strains was 72, which was the *pks^-^* strain **(Fig. 5A).** Furthermore, there were similar numbers of *pks^+^* strains from UCs (median: 45, IQR: 43-47) and other strains harboring *pks* island (median: 44, IQR: 43-46, *P*=0.816, Mann □ Whitney *U* test), while there was a significant difference in *pks^-^ strains* (median: 48, IQR: 46.5-55, *P*=0.003, Mann □ Whitney *U* test, **Fig 5B**). For the ARGs, the macrolide resistance gene *mphA* (0.00% in UC patient *pks^+^* strains *vs.* 27.78% in *pks^-^* strains; *P*=0.045, chi-square test), the diaminopyrimidine resistance gene *dfrA17* (0.00% in UC patient *pks^+^* strains *vs.* 30.56% in *pks^-^* strains; *P*=0.024, chi-square test), and the multidrug resistance gene *mdtM* (23.07% in UC patient *pks^+^* strains *vs.* 58.33% in *pks^-^* strains; *P*=0.029, chi-square test) were significantly enriched in the *pks^-^* strains **(Fig. 5C, Table S5).**

**Fig. 5.**
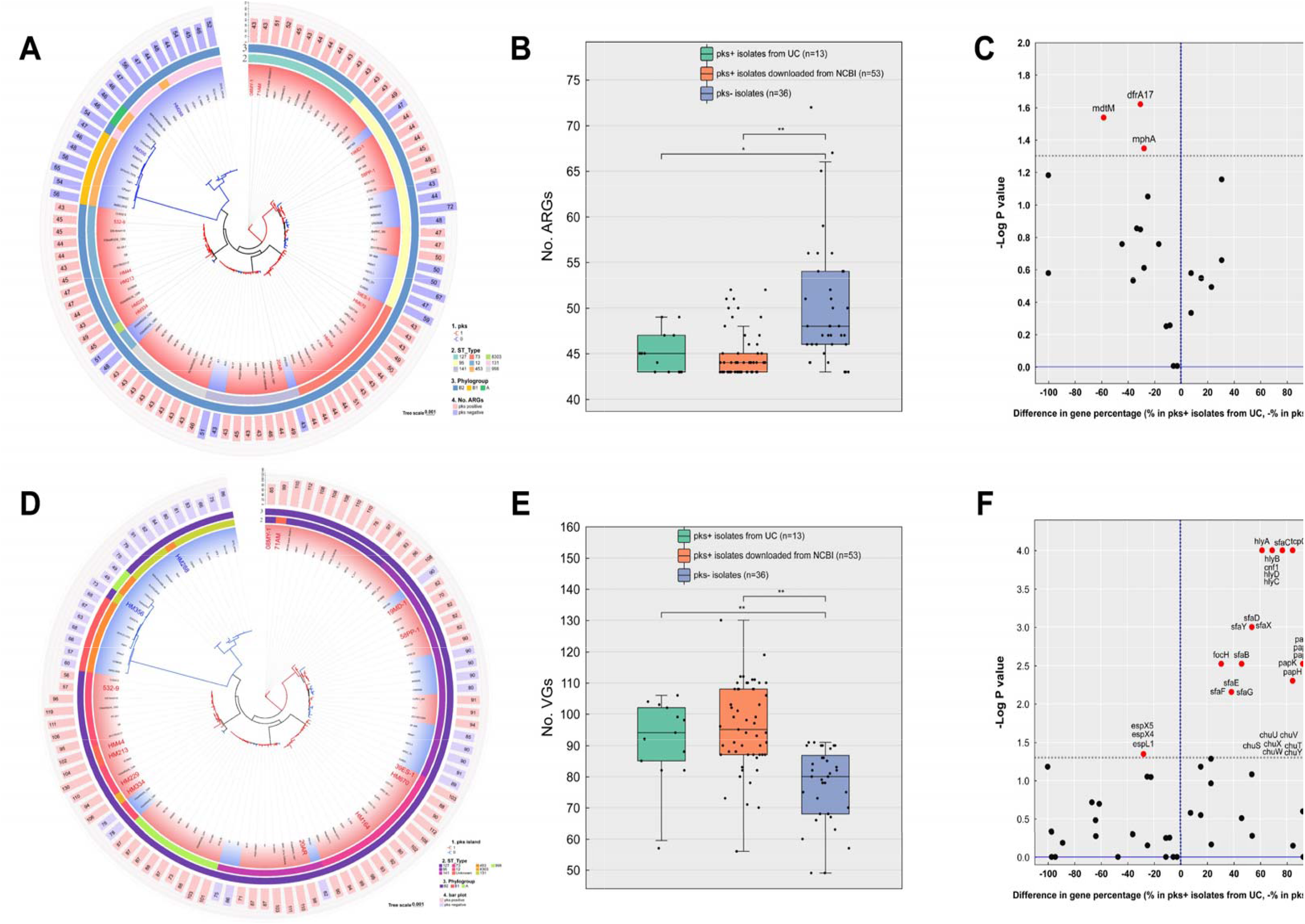
Virulome and resistome profiling of UC patient *pks^+^ E. coli* strains, other *pks^+^* strains and *pks^-^* strains. The results were obtained from 102 selected *E. coli* strains. A: the core gene-based phylogenetic tree with the number of ARGs; B: the box plot of the ARGs, * *P* value <0.05, ** *P* value <0.001; C: the difference in ARGs between the UC patient *pks*-positive *E. coli* strains (n=13) and the *pks*-negative *E. coli* strains (n=36), the dashed line indicates *P* value < 0.05; D: the core gene-based phylogenetic tree with the number of VGs; E: the box plot of the VGs, * *P* value <0.05, ** *P* value <0.001; F: the difference in ARGs between the UC patient *pks*-positive *E. coli* strains (n=13) and the *pks-*negative *E. coli* strains (n=36), the dashed line indicates *P* value < 0.05.

For VGs, the maximum quantity was 130, which was associated with the *pks^+^* strain **(Fig. 5D)**. The two *pks^-^* strains belonging to ST453 and phylogroup A possessed the lowest number of VGs (49). Contrary to the quantity of ARGs, the VGs of the *pks^+^* strains including strains from UC patients (median: 94, IQR:83.5-102.5) and other source (median:99, IQR:95, IQR:87-108) were significantly more than that of *pks^-^*strains (median: 80, IQR:80-88.25, *P*=0.001 and *P*<0.001, Mann □ Whitney *U* test, **Fig. 5E**). Among the VGs, the *pks^+^* strains from UC patients presented significantly greater percentages of VGs encoding adherence factors (*papC*, *papD*, *papF*, *papH*, *papJ*, *papK*, *sfaB*, *sfaC*, *sfaD*, *sfaE*, *sfaF*, *sfaG*, *sfaX,* and *sfaY*); nutritional/metabolic *factors (ChuA*, *chuS*, *chuT*, *chuU*, *chuV*, *chuW*, *chuX,* and *chuY*); the effector delivery system (*vat*); invasion factor (*aslA*); exotoxin (*hlyA*, *hlyB*, *hlyC*, *hlyD,* and *cnf1*); and immune modulation factor (*tcpC*). Moreover, the percentages of effector delivery system genes *espX4*, *espX5,* and *espL1* in the *pks^-^* strains were greater than those in the *pks^+^* strains from the UC patients (27.78% *vs.* 0.00%, *P*=0.045; chi-square test; **Fig. 5F**; **Table S4**).

We also compared the differences between ARGs and VGs among *pks^+^* strains from UC patients, UTI patients, and healthy individuals. For the ARGs, the quantity of *pks^+^* strains from UC patients, UTI patients, and healthy individuals were not significantly different (*P*=0.693 and 0.964; **Fig. S2A**). For VGs, the quantity of *pks^+^* strains from UC patients was not significantly different from that from UTI patients or healthy people (*P*=0.759 and 0.316, **Fig. S2B**). Similarly, regarding the percentage of ARGs, there was no significant difference in ARG abundance between the *pks^+^* strains from UC patients and those from UTI patients, and the percentages of the ARGs *ANT (3’’)-IIa* and *tet*(B) were greater for the *pks*^+^ strains from UC patients than for those from healthy people (**Fig. S2C, D**). Although various kinds of VGs were differentially expressed between the UC group and the healthy people group, only *foc*D (adherence factor, higher in healthy people), *tcp*C (immune modulation factor) and *pap*B (adherence factor) were significantly different **(P<0.05; Fig. S2E**), and two VGs, *pap*L (adherence factor) and *iut*A (nutritional/metabolic factor), were significantly different between the UC group and the UTI group (*P*<0.05; **Fig. S2F**).

### Bacteriophages and Genomic Islands in HM229 and other strains of *E. coli*

Subsequently, we compared only one complete genome of HM229 from UC patients with 7 other selected downloaded complete genomes from different phylogroups, including the reference strain IHE3034 (**Table 3 and Table S5**). The complete genome of HM229 was annotated to a 5028,419 bp long chromosome and a 122,249 bp long plasmid compared with the reference strain IHE3034, which has a 5108,383 bp chromosome size and no plasmid (**Fig. 6**). In the gene map of HM229, we identified all the predicted genomic islands (GI), and the *pks* island was coded G15 (located at 2048, 908-2104, 007 bp). The reference isolate IHE3034 contained 49 GIs, and *the pks island was* also identified in the genome map coded GI24. Moreover, 7 bacteriophages (labeled P1-P7) were identified in the sequence of HM229, while the reference isolate IHE3034 was used to filter out 12 bacteriophages. According to the genome map, no bacteriophage was present in the peripheral region of the pks island (**Fig. 6**). Moreover, the size of the chromosome, number of CDSs, and number of genes encoding tRNAs varied between HM229 and the other 7 selected strains (Table 3). The genome maps of all 8 strains are shown in **Fig. S3.**

**Fig. 6.**
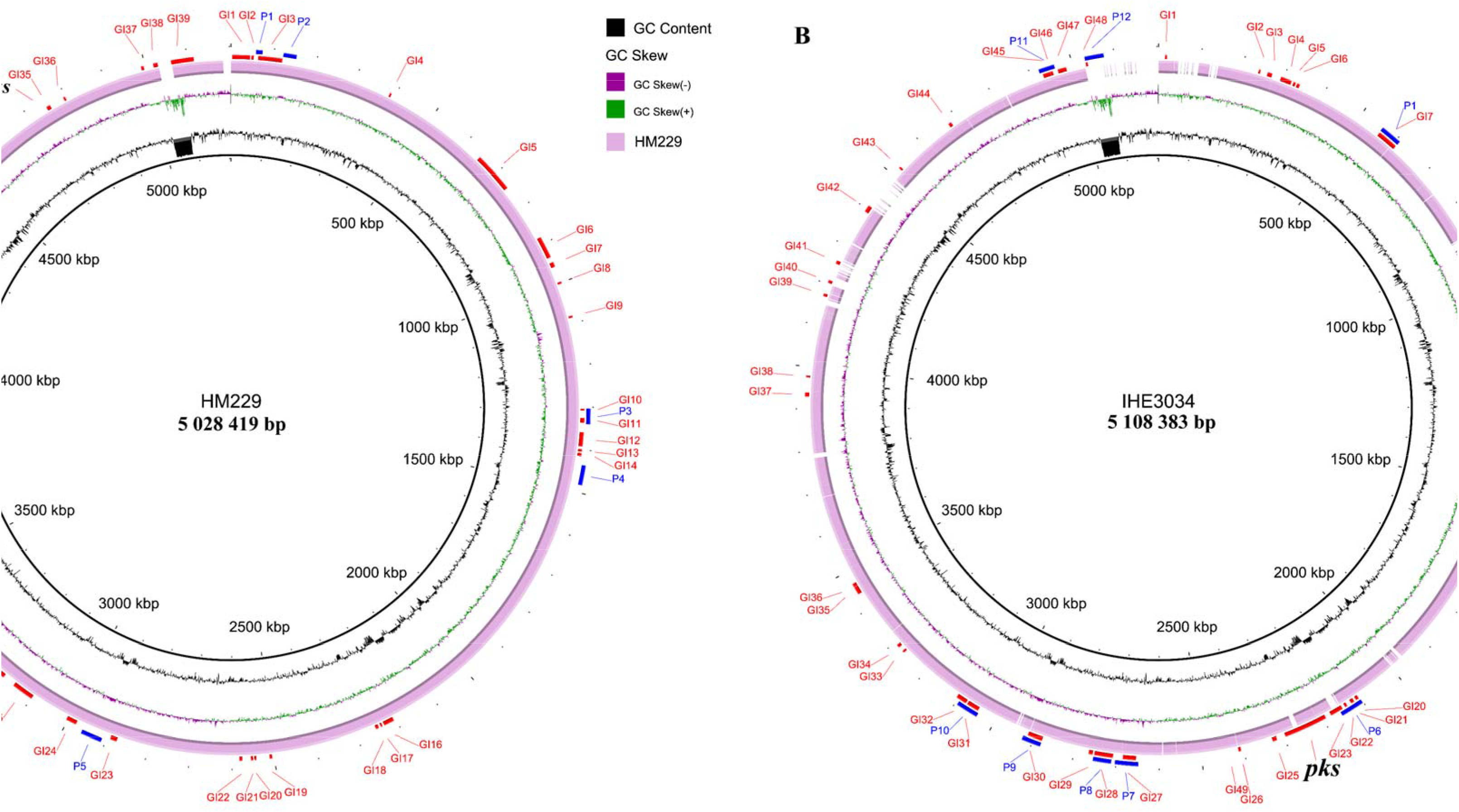
A circular genome map of HM229 cells isolated from UC patients (A) and the reference isolate IHE3034 (B). In the two maps, P represents bacteriophage or bacteriophage regions, and GI represents the genomic island.

**Table 3:**
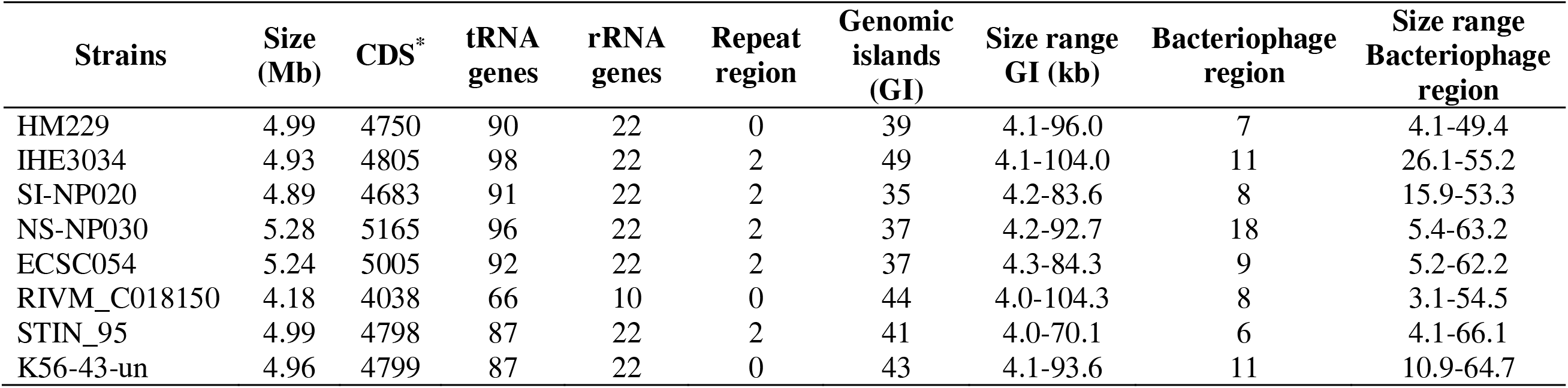
Comparison of basic features of the HM229 chromosome and 7 selected strains.

The plasmid encoding the 148 CDS was constructed for HM229. In all CDSs, 6 IS family transposases (IS1, IS3, IS5, IS6, IS21, and IS91) were identified, indicating the high dynamics of the plasmid or the components of the plasmid. Moreover, there are toxic genes or virulence genes for *rel*E, *pem*K, *esp*C, *pnd*A and *ylp*A in the plasmid; these genes include serine proteases with enterotoxic and cytotoxic activities and type I and type II toxin-antitoxin (TA) systems. Additionally, several antimicrobial genes were also identified, including bla, which encodes the beta-lactamase SHV-1; *tet*A, which encodes a tetracycline resistance protein; *emr*E, which encodes a multidrug efflux protein; and *ant*1, which encodes spectinomycin 9-adenylyltransferase (**Fig. 7**). In addition to those of HM229, the genomes of ECSC054 and STIN_95 also contained one plasmid, and the isolate NS_NP030 contained 3 plasmids (coded P1-P3). We also used BRIG to compare the six plasmids, and the figure shows partial homology between the HM229 plasmid and the ECSC054 plasmid, between the STIN_95 plasmid, and NS_NP030_P1. The residual plasmids NS_NP030_P2 and NS_NP030_P3 had no homology with the aforementioned plasmids (**Fig. S4**).

**Fig. 7.**
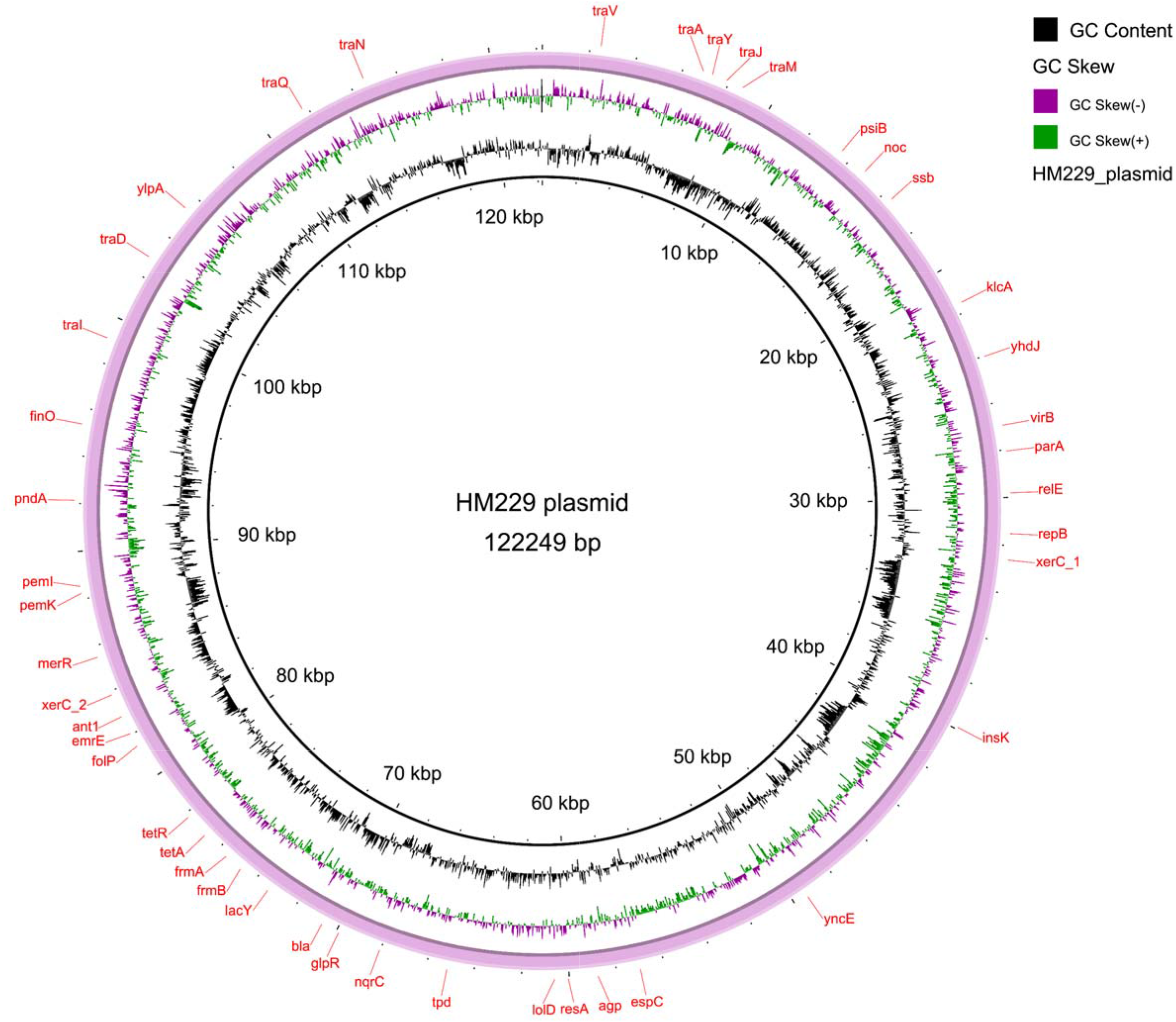
A circular map of the plasmids used to isolate HM229 cells from UC patients.

### Conserved genomic blocks in the chromosome of HM229

All 8 selected *E. coli* strains shared a common core genome of approximately 3.54 Mb organized into linear conserved blocks (LCBs). Except for strain RIVM_C018150, the results of this comparative alignment showed that the chromosomes of the compared strains were organized into 20 LCBs (**Fig. 8**). Overall, the LCB of HM229 was identical to that of 4 of the 7 selected strains, including the strain ECSC054 from phylogroup D, highlighting the conserved genomic skeleton of the *pks*^+^ *E. coli* strains. The *pks* island was in LCB 6, although the location of LCB 6 in the genome was a translocation in some strains, such as NS_NP030, RIVM_C018150, and SI-NP020. NS_NP030 and SI-NP020 were isolated from bovines and belonged to phylogroups B1 and A, respectively. RIVM_C018150 was a clinical sample and belonged to phylogroup F. These different locations, which significantly changed in arrangement from LCB (NS_NP030, RIVM_C018150), indicated that *pks* island introduction might have significant recombination in the genomic skeleton of nonphylogroup B2 strains.

**Fig. 8.**
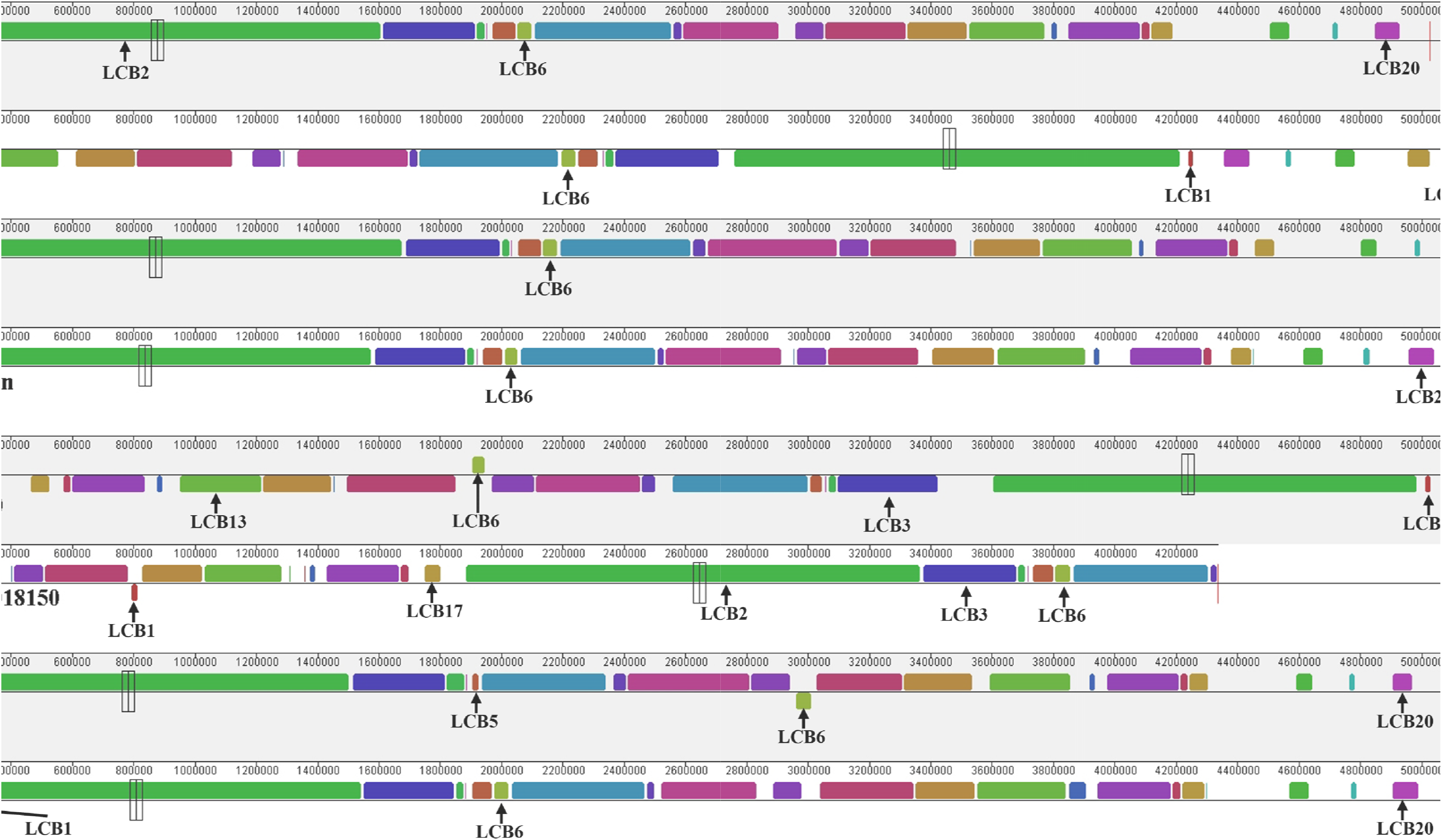
Whole-genome alignments of the species created by Mauve. The colored rectangles represent LCBs. The sizes of the rectangles are proportional to the genomic extensions of the LCBs. The isolate HM229 was used as a reference, and the LCBs were ordered according to the reference.

### Conserved *PKS* Island in HM229 Cells

Although the isolate HM229 and the reference isolate IHE3034 belonged to different STs, the *pks* island genome structure of HM229 was identical to that of the reference isolate, including the periphery of tRNA-Asn, the bacteriophage integrase gene *intP4*, the insertion sequence (*IS*1400 family) and the putative transposase gene (**Fig. 9**). The identical genome structure of these strains indicated that the *pks* island of HM229 could have a function equal to that of IHE3034. A comparison of the *pks* island genome structure among the 8 selected strains revealed that most of the genes encoding the biosynthetic machinery on the *pks* island (19 genes from *clb*A to -S) were highly conserved, except for the three genes *clb*B, *clb*J, and *clb*K. However, the peripheral elements of the aforementioned 19 genes were variant; for example, the insertion sequence of 4 out of the 8 strains belonged to the *IS*3 family; two isolates, SI-NP020 and NS_NP030, had tRNA-Asn, an integrase gene and a transposase gene deletion; and three strains, ECSC054, C018150 and STQIN_95, added to the *yee*O gene, which encodes the toxic metabolite efflux MATE transporter upstream and changed into the IdtA gene, which encodes the L, D-transpeptidase downstream. The clbK-J fusion was found in two strains, K56_43_un and NS_NP030, and these two strains belonged to phylogroups B2 and B1, respectively, indicating that the *clb*K-J fusion is likely to occur in all phylogroups without known proportions. In addition, the clbB gene was truncated in phylogroups A and B1 but not in B2. Another feature of the 4 strains that belonged to phylogroup B2 according to the genome structure analysis was that all the strains had tRNA-Asn, indicating that active protein synthesis occurs in *pks*-positive strains belonging to B2.

**Fig. 9.**
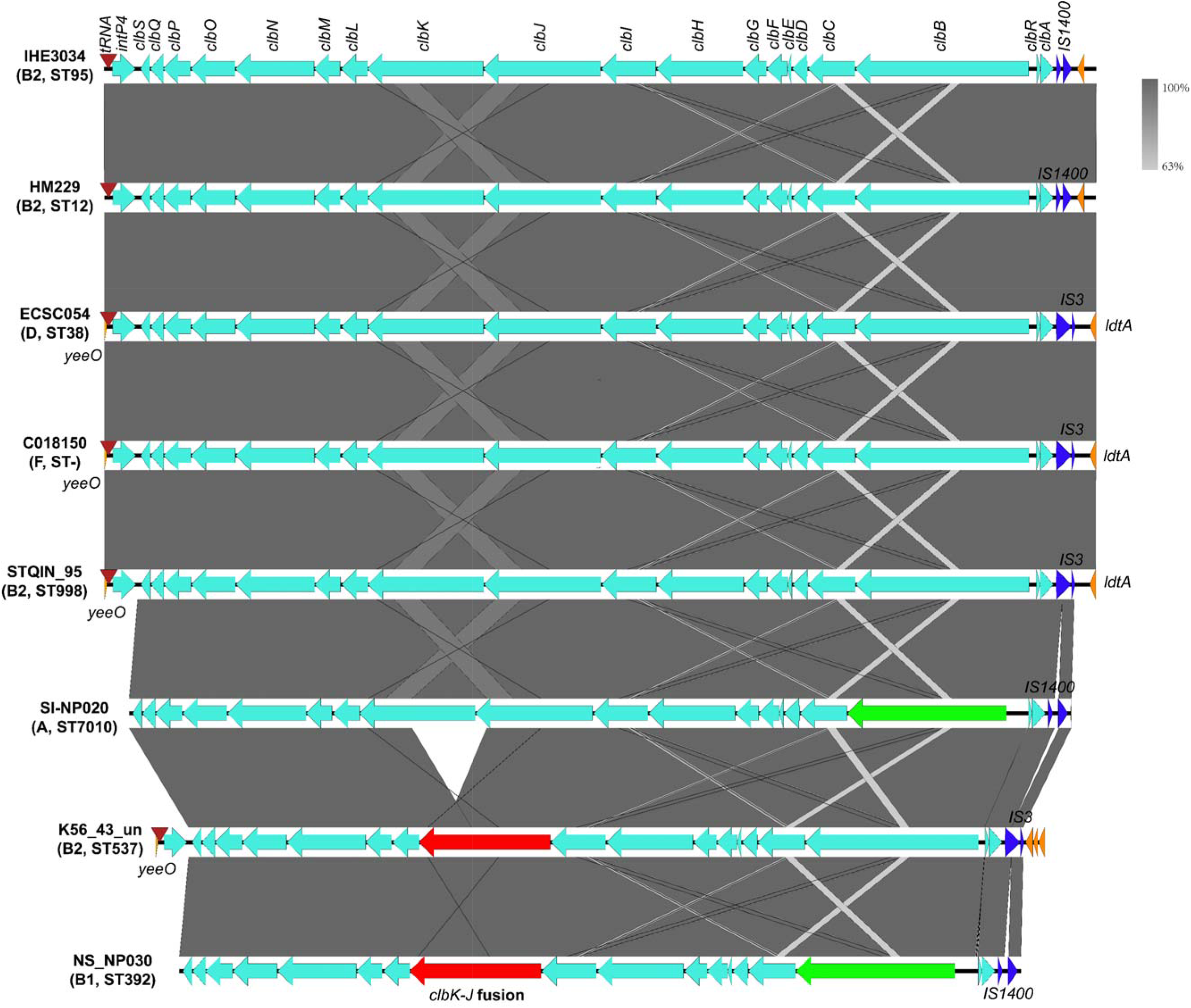
The *pks* island structure of strains belonging to different phylogroups. IHE3034 was the reference isolate, HM229 was the isolate from UC patients, and ECSC054, C018150, SI-NP020, and NS_NP030 were the strains belonging to phylogroups D, F, A, and B1, respectively. Compared with those of HM229, the clades of STQIN_95 and k56_43_un, which belong to the B2 phylogroup, were different.

### Single nucleotide polymorphisms (SNPs) in *pks* island

The snippy program identified SNPs in the 8 selected strain genomes and *pks* island sequences. There were 2722 core SNPs among the 8 strains, and the number of core SNPs among the 8 *pks* island sequences was 0. When comparing HM229 and the reference isolate IHE3034, 31030 SNP sites were identified, and only 8 SNP sites were screened in the *pks* island sequence. The SNP sites on the *pks* island of SI-NP020 (phylogroup A) and NS-NP030 (phylogroup B1) were 82 and 94, respectively, which were significantly greater than those of the other strains. Interestingly, the two strains were isolated from bovines.

## Discussion

Colibactin, which is produced by *pks* islands and identified in specific Enterobacteriaceae members, has emerged as an essential virulence factor implicated in the *progression* of CRC, meningitis, and septicaemia [12]. Many studies in the past have reported the involvement of colibactin in CRC, *as it plays* an important role *in the interaction* between host cells and *the* microbiota *during* the progression of CRC, *suggesting that* colibactin *is* an important virulence factor *that demands* prompt attention due to its far-reaching implications [50].

In the *present* study, whole*-*genome-wide *comparisons* of *E. coli* isolates from in-house culture collection and from a public database *were performed to obtain* insight into island acquisition and evolution. The in-house genome collection, derived from human patients (n=15), revealed that 13 of these genomes *harbored* the *pks* island. The score of the study was further broadened by including 2654 *E. coli* genomes from *the* NCBI database, unveiling the presence of *pks islands* in 158 isolates. The subsequent phylogenomic analysis *revealed* that 13 *pks*-positive isolates from in-house culture and 158 genomes from the public database belonged to phylogroup B2. This aligns seamlessly with earlier research findings [28, 51–53]. The 171 genomes exhibiting *pks* positivity *were subjected to* comprehensive in silico typing techniques to discern distribution patterns across diverse *E. coli* subtypes. Notably, the majority of the 13 *pks*^+^ isolates were identified as belonging to ST12 (30.77%, 4/13) *or* ST73 (23.07%, 3/13). Conversely, the 158 *pks*^+^ isolates sourced from NCBI *showed* dominance in ST73 (29.75%, 47/158), ST95 (22.78%, 36/158), and ST127 (15.19%, 24/158). These findings align consistently with outcomes reported in earlier investigations. We meticulously crafted a phylogenetic tree to *determine* the circumstances surrounding the acquisition of *pks* sequences by *pks*^+^ *E. coli*. Our findings illuminated a clustering of the core genome primarily within lineages of *the* CC12, CC14, CC95, and CC73 *clonal complexes*. This *finding* supports the hypothesis that the introduction of the *pks* island into CC12, CC14, CC73, and CC95 occurred through horizontal acquisition by their most recent common ancestor, subsequently followed by vertical transmission with gradual *pks* divergence over time [54]. Notably, the analysis of the phylogenetic tree *revealed* that CC73 *exhibited* the longest divergence time (91.97), while CC95 *had* the earliest average divergence time (26.23), indicating that CC95 was the first to acquire the *pks* island. *Furthermore, a pangenome* analysis of *pks*+ *E. coli* strains *revealed* that *the* core genome size *was not significantly different* between UC patients and *those in* the downloaded NCBI dataset. The heatmap shows that the *pks* island genes *clb*J, *clb*H, *clb*M, and *clb*I are missing from *pks*^+^ *E. coli* strains from UC patients, except for the HM229 strain. The absence of these genes may be due to genomic deletion or mutation. Random mutations or events such as recombination can lead to the loss of specific gene sequences.

Whole-genome-based virulome and resistome analyses revealed that 102 *E. coli* strains contained 72 antibiotic resistance genes among the *pks*-strains and 130 virulence genes among the *pks*^+^ strains (Supplementary Table S4). On the basis of our study, it was found that, compared with pks-negative isolates, *pks*^+^ isolates contain fewer antibiotic resistance genes and a greater number of virulence genes. Our results are in line with those of previous studies, which showed low antibiotic resistance and virulence in *pks*^+^ isolates [55, 56]. Among the virulence genes identified in the *pks*^+^ isolates, a significantly higher number of genes were associated with adherence factor, invasion factor, exotoxin, and immune modulation factor. The large number of virulence genes identified in the *pks*^+^ isolates is consistent with the findings of previous reports based on PCR-based observations of bacteremia isolates [52]. The comparative genomic alignment of 8 strains revealed 20 linear conservative blocks (LCBs), with HM229 having an identical orientation to 4 of the 7 strains, highlighting the conserved genomic skeleton of the *pks*^+^ isolates. The high conservation of the *pks* island suggested that colibactin is an important genotoxin.

## Conclusion

The prevalence of colibactin-producing *E. coli* isolates was found to be very high in UC patients, and most of the *pks*^+^ isolates belonged to Phylogroup B2. The *pks* island distribution pattern supports the diagnostic system and helps to understand the clinical implications of the genotoxic nature of *pks*^+^ isolates. The *pks* island phylogeny indicates that the *pks* island spread through horizontal gene transfer. The *pks*^+^ isolates demonstrated high virulence and low antibiotic resistance.

## Data Availability

Data will be made available upon request.

## Acknowledgment

We acknowledge funding from R01 CA231283 to support this study.

---

**Fig. S1.**
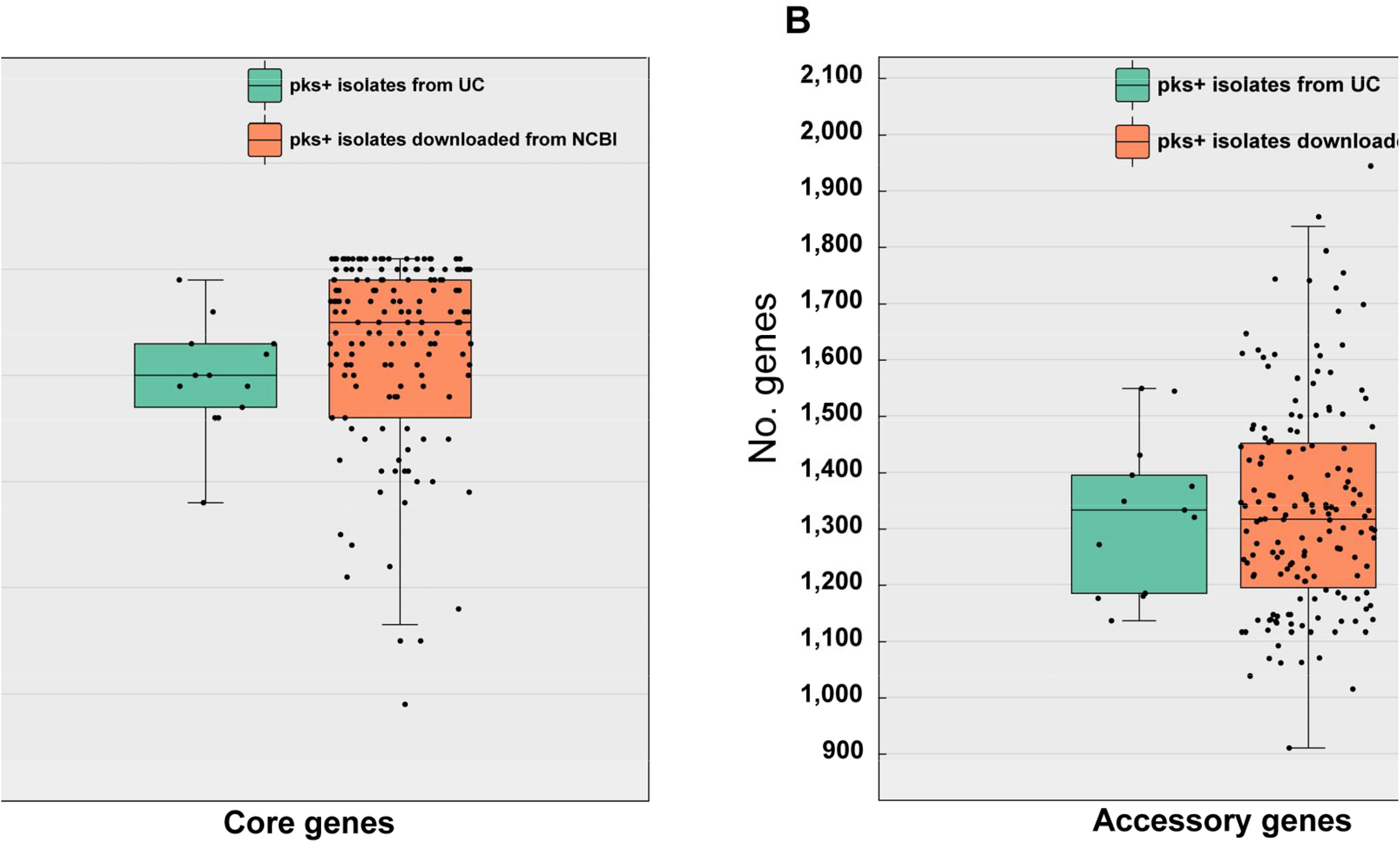
Total number of core and accessory genes between the *pks*-positive strains from UC patients and the other isolates downloaded from the NCBI. A. Total number of core genes. B. Accessory genes. The data are shown as box plots, and the horizontal box lines represent the first, median, and third quartiles. Whiskers denote the range of points within the first quartile – 1.5× the interquartile range and the third quartile + 1.5× the interquartile range. Significant levels are labeled with an asterisk using the Mann–Whitney U test.

**Fig. S2.**
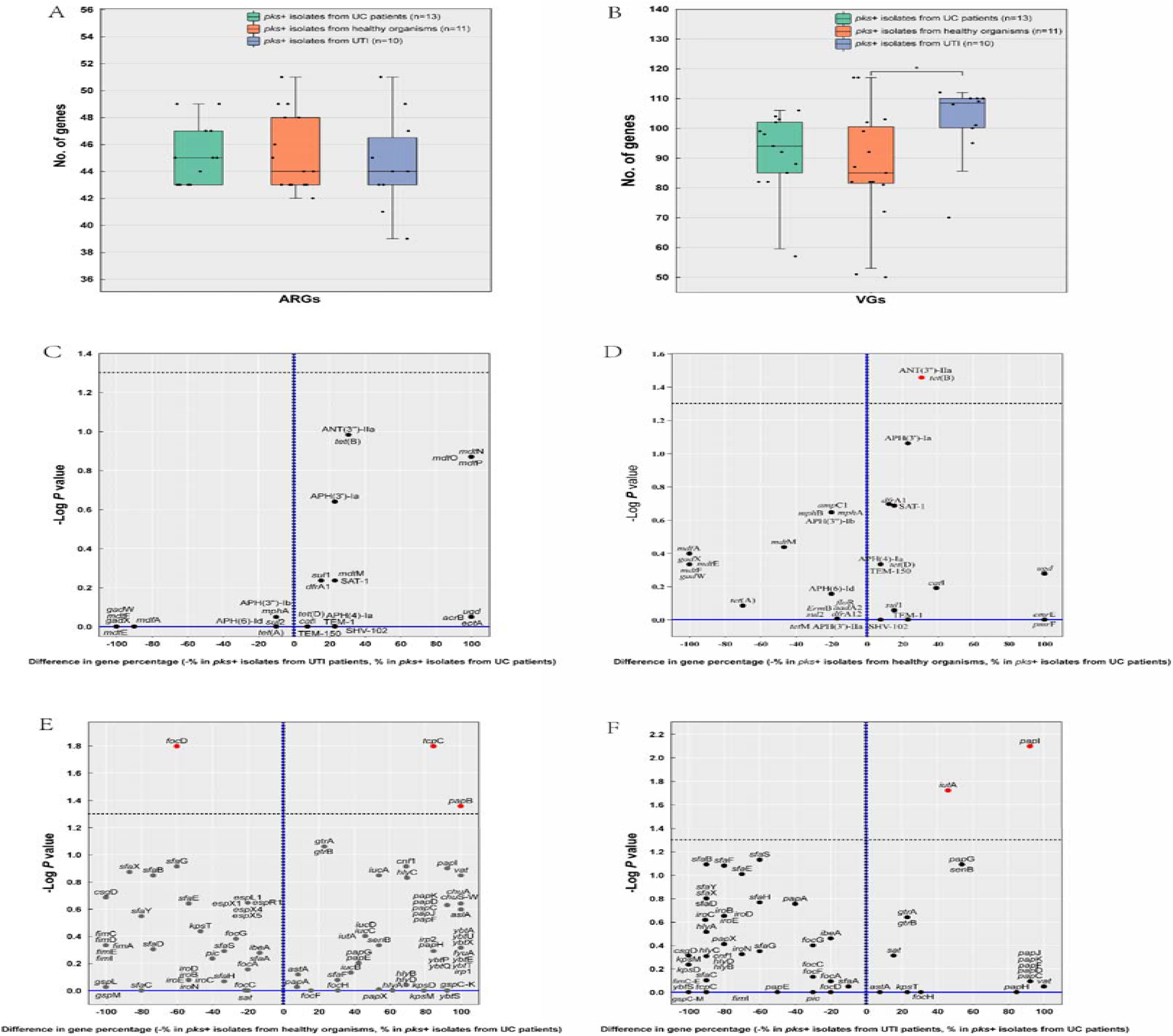
Virulome and resistome profiles of UC patient *pks^+^ E. coli* strains (n=13), UTI patient *pks^+^* strains (n=10) and healthy organism *pks^+^* strains (n=15). A: Box plot of ARGs; B: Box plot of VGs; * P value <0.05; C: Difference in ARGs between the UC patient *pks^+^ E. coli* strains (n=13) and UTI patient *pks^+^* strains (n=10); the dashed line was *P* value < 0.05; D: Difference in ARGs between the UC patient *pks^+^ E. coli* strains (n=13) and healthy organism *pks^+^* strains (n=10), the dashed line was *P* value < 0.05; E: Difference in VGs between the UC patient *pks^+^ E. coli* strains (n=13) and UTI patient *pks^+^* strains (n=10), the dashed line was *P* value < 0.05; F: Difference in VGs between the UC patient *pks^+^ E. coli* strains (n=13) and healthy organism *pks^+^* strains (n=10), the dashed line was *P* value < 0.05.

**Fig. S3.**
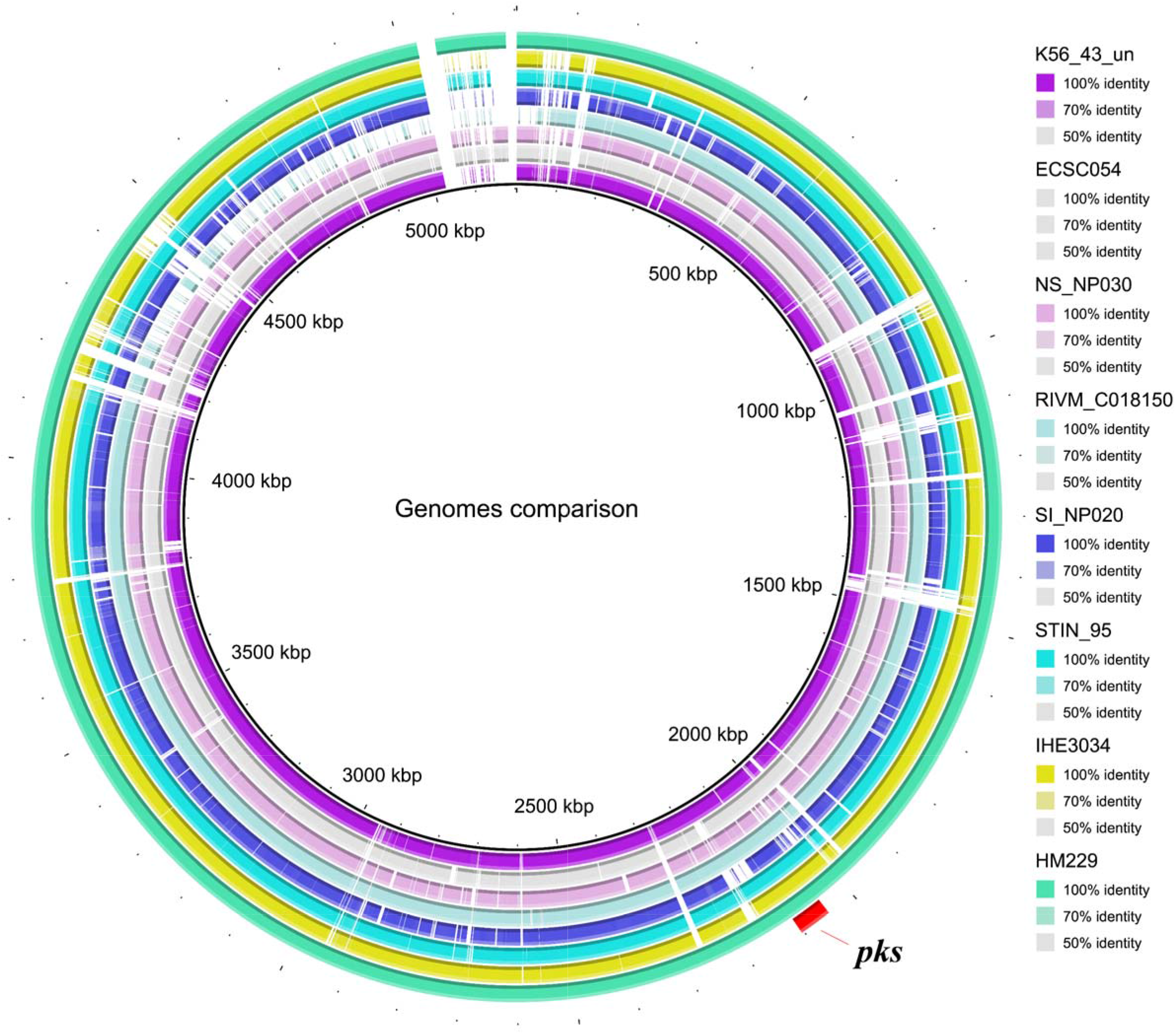
Circular genome map of 8 selected strains. HM229 was used as a reference.

**Fig. S4.**
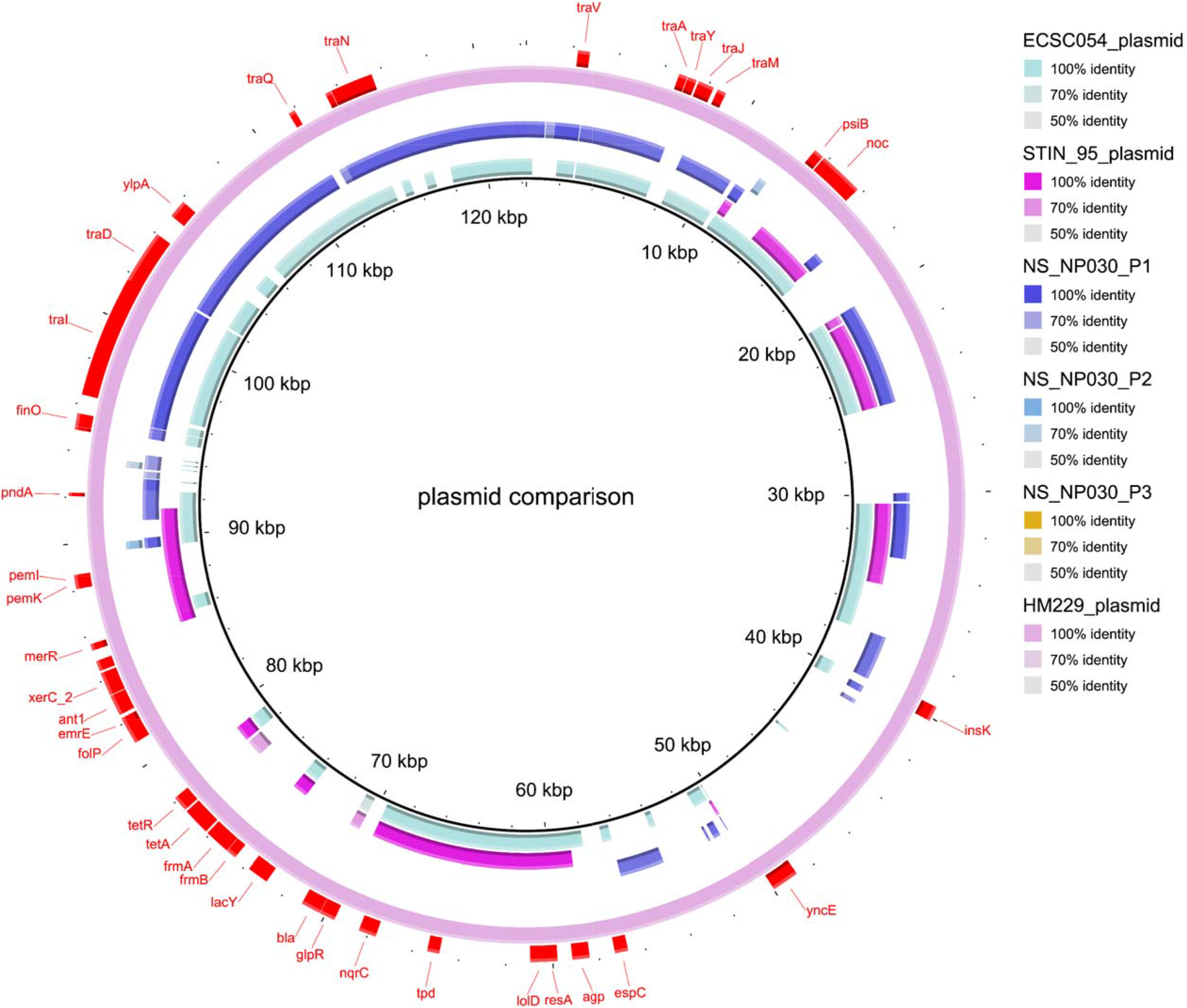
Comparison map of plasmids from the 8 selected strains. The plasmid HM229 (HM229_plasmid) was used as a reference. The strains HM229, ECSC054, and STIN_95 each contained one plasmid encoding the HM229 plasmid, the ECSC054 plasmid, and the STIN_95 plasmid. The isolate NS_NP030 contained 3 plasmids and encoded NS_NP030_P1, NS_NP030_P2, and NS_NP030_P3.

**Supplementary Table S1:** Details of the isolates obtained from 15 patients with ulcerative colitis and additional isolates downloaded from the NCBI database.

**Supplementary Table S2:** Serotypes of 171 pks-positive isolates

**Supplementary Table S3:** In-depth analysis of the virulome and resistome profiles of the 102 *E. coli* isolates.

**Supplementary Table S4:** The percentages of virulence genes (VGs) and antibiotic resistance genes (ARGs) in *pks*^+^ isolates from patients with ulcerative colitis (UC) and *pks*^-^ isolates.

**Supplementary Table S5:** Eight selected isolates used for genome mapping and SNP analysis.

